# Transplantation of human iPSC-derived retinal cells in the cone-dominant Northern tree shrew

**DOI:** 10.64898/2026.10.28.761748

**Authors:** Ching Tzu Yu, Sangeetha Kandoi, Hannah M. Follett, Eeshika Mitra, Ramesh Periasamy, Kenneth P. Allen, Phyllis Summerfelt, Susan Freling, Daniel M. Lipinski, Joseph Carroll, Deepak A. Lamba

**Affiliations:** Cell Biology, Neurobiology & Anatomy, Medical College of Wisconsin, Milwaukee, WI, USA; Department of Ophthalmology, University of California San Francisco, San Francisco, CA, USA; Solomon H. Snyder Department of Neuroscience, Johns Hopkins University, Baltimore, MD, USA; Ophthalmology, Genentech, South San Francisco, CA, USA; Ophthalmology & Visual Sciences, Medical College of Wisconsin, Milwaukee, WI, USA; Department of Microbiology and Immunology, Biomedical Resource Center, Medical College of Wisconsin, Milwaukee, WI, USA; Max Planck Florida Institute for Neuroscience, Jupiter, FL, USA

**Author notes:** **Correspondence:** Joseph Carroll, Deepak A. Lamba. These authors contributed equally (Ching Tzu Yu, Sangeetha Kandoi, Hannah M. Follett).

**Keywords:** Northern tree shrew, Multimodal in vivo imaging, AOSLO, retinal cell transplantation, species-specific immunosuppression

## Abstract

**Purpose:** To characterize the transplantation and survival of photoreceptors generated from human induced pluripotent stem cell (hiPSC)-derived retinal organoids in the Northern tree shrew (*Tupaia belangeri*, TS).

**Methods:** TS eyes were injected with retinal cells dissociated from GFP-expressing hiPSC-derived retinal organoids (12 animals; n=22 eyes). 15 eyes had prior damage induced by either experimental retinal detachment (n=11) or ATP-induced retinal damage (n=4), and seven eyes were undamaged. Retinal organoids were treated with Notch pathway inhibitor 2 days before transplant. TS retinal structure and cellular details were assessed post-transplantation for up to 8 weeks using scanning laser ophthalmoscopy (SLO), optical coherence tomography (OCT), and adaptive optics SLO (AOSLO). Histological analysis was done at the end of the experiment.

**Results:** Cell transplants in eyes treated with a simple immunosuppression treatment failed to survive for more than 2 weeks (n=10 eyes). Robust cell survival was observed in the eyes treated with a complex immunosuppression cocktail for up to 8 weeks (n=12 eyes). The transplanted cells were confirmed to be subretinally located using OCT imaging and the cell fate was confirmed to be human photoreceptors by histology. Finally, AOSLO imaging captured structures resembling photoreceptor inner segments.

**Conclusions:** Our results demonstrate survival of transplanted cells using a species-specific immunosuppression treatment. Future studies should aim to explore the long-term survival and function of the transplanted cells.

## Introduction

Photoreceptor transplants have been extensively explored as therapeutic interventions to replace cells in end-stage retinal degeneration in several pre-clinical animal models, including non-human primates (NHP), dogs, mice, rats, and 13-lined ground squirrels (13-LGS) (Ripolles-Garcia et al., 2022, Aboualizadeh et al., 2020, Zhu et al., 2017, Yu et al., 2024). Aside from the main difference of the animal model, many other parameters can also influence the outcome of the post-transplant result. In xenogeneic versus allogeneic transplant studies, comparing outcomes allows us to understand how donor and host cells interact across species or within the same species, respectively (Ho et al., 2024). To date, several experimental parameters have been evaluated aside from the animal model and origin of transplant to improve the outcome of donor cell survival, reformation of synaptic connectivity, and restoration of visual function in the host (Uyama et al., 2022, Zhu et al., 2017, McLelland et al., 2018, Pearson et al., 2012). A few other considerations, including immunosuppression protocols, transplant delivery route, the form of the transplanted material (i.e., scaffolds, dissociated cells), and the selection of transplanted cell types (i.e. mixed cell type, isolation of photoreceptors), have also been adjusted to optimize the outcome in numerous animal models (Ripolles-Garcia et al., 2022, Gagliardi et al., 2018, Ludwig et al., 2019). Nevertheless, there has been no one standard combinatorial approach to transplantation parameters that can be applied across all species. Therefore, optimizing the transplantation parameters across numerous novel animal model systems is imperative, as it would advance our understanding of photoreceptor transplants prior to getting closer to clinical trials.

The Northern tree shrew (*Tupaia belangeri*) is a valuable animal model, particularly in vision research (Savier et al., 2021, Norton et al., 2022). It has been used as an alternative to NHP models in numerous applications, including visual field assessments and high-resolution non-invasive retinal structure and functional imaging (Xiao et al., 2017, Van Hooser et al., 2013, Sajdak et al., 2019, Gaffney et al., 2021). Historically, they have proven to be a valuable research model for studying myopia,(Ward et al., 2016, Sherman et al., 1977, Abbott et al., 2011, Khanal et al., 2023) glaucoma, and diabetic retinopathy (Samuels et al., 2018, Gorbatyuk et al., 2022). One particularly attractive characteristic of the tree shrew is their cone-dominant retina, where up to 95% of photoreceptors are cones (Müller and Peichl, 1989). Here, we sought to optimize the transplant conditions for the delivery of human-induced pluripotent stem cell (hiPSC)-derived retinal cells in the tree shrew retina. We further explored how these cells survive in a degenerated cone dominant host retina. The findings of this study demonstrate survival of transplanted hiPSC-derived retinal cells in both damaged and undamaged tree shrew retinae through multimodal *in vivo* imaging and histology for up to 8 weeks post-transplantation.

## Materials and Methods

### Animals

The northern tree shrews (*Tupaia belangeri*) used in this study were obtained from Max Planck Florida Institute of Neuroscience Center, Jupiter, FL, USA. All experiments were performed with the approval of the Institutional Animal Care and Use Committee of the Medical College of Wisconsin (MCW) and in accordance with the ARVO Statement for the Use of Animals in Ophthalmic and Vision Research. 12 tree shrews were used, and 22 eyes were injected with dissociated cells from human iPSC-derived retinal organoids. For all procedures, tree shrews were initially anesthetized with 4-5% inhaled isoflurane, mixed with 100% oxygen, using an induction chamber attached to an anesthetic machine (VetEquip Inc., Livermore, CA, USA). Once the animal was anesthetized, it was removed from the chamber and subcutaneously infused with dexmedetomidine (0.3 mg/ kg/ hr) mixed with normal saline. Supplemental anesthesia was administered through a mask and 1-2% inhaled isoflurane to maintain the sedation state of the animals throughout the sessions. Some animals received subcutaneous injections of atropine (0.3 mg/kg) at the beginning of the session to control excessive salivation. At the end of the session, the animals were recovered from sedation via subcutaneous injection of atipamezole (0.5 mg) to reverse the effects of dexmedetomidine. Throughout the session, the animal’s breathing rate was monitored and recorded every 15 minutes. The body temperature of the animal was maintained at 37°C by placing the electrical heating pad in the animal imaging cassette.

### Immunosuppression treatment

For the first 10 eyes undergoing cell transplantation, the immunosuppression protocol was adapted from our previous 13-LGS study (Yu et al., 2024). The animals were treated with cyclosporine A (100 mg/L) in drinking water for 1 week prior to transplantation and continued until the end of the experiment. For the remaining 12 eyes, a species-specific immunosuppression protocol was further modified. In this protocol, tree shrews were subjected to immunosuppression treatments for 1 week prior to transplantation and continued until the end of the experiment. Weekly, the animals were provided with methylprednisolone (10 mg/kg) via intraperitoneal injection and doxycycline (10 mg/kg) mixed in applesauce. Daily, the animals were treated with sirolimus (1 mg/kg) and azathioprine (10 mg/kg) mixed into their applesauce and cyclosporin A in their drinking water. Furthermore, the animals were treated with basiliximab (0.3 mg/kg) through intravenous injection on the day of transplantation and four days post-transplantation. The weight of the animal was recorded at the start of the immunosuppression day and during each imaging session until the end of the experiment.

### Induced retinal degeneration in tree shrews

Before each surgical session, the animals were anesthetized, and the eyes were dilated and cyclopleged as described in the retinal imaging section of the methods. Following the animal preparation, the eye was topically sterilized with betadine (Alcon, Fort Worth, TX, USA) on the surface and around the regions near the eye. The eye was coated with either Gonak solution (Sigma Pharmaceuticals, North Liberty, IA, USA) or Celluvisc (Allergan Inc., Markham, ON, Canada). A small sterile circular coverslip was placed against the cornea to obtain visualization of the retina. The retina was visualized via a Leica Proveo 8 ophthalmic microscope equipped with intraoperative optical coherence tomography (OCT) (Leica Biosystems, Wetzlar, Germany) throughout the injection procedure. In both damage models, a 2 - 3 mm retinal region directly nasal to the circular optic disc was targeted for inducing damage.

Chemically-induced photoreceptor degeneration was created in 4 eyes via 10 μL intravitreal injections of ATP (Thermo Fisher Scientific, Waltham, MA, USA) with concentrations ranging from 0.0075-0.01 M, depending on the vitreous volume and axial length of each eye. The ATP solution was delivered using a 1 mL insulin syringe with the needle tip placed directly above the targeted retinal region, nasal to the optic nerve. For the remaining 11 eyes, retinas were mechanically detached to induce photoreceptor degeneration. A 25-gauge trocar was first used to make a small cut 1 mm from the limbus of the eye, and a 100 µL Hamilton syringe with a 33-gauge blunt needle was used to inject 25-100 µL of 0.9% normal saline (Phoenix, Manhattan, KS, USA) delivered subretinally to induce detachment. Following the injections, the eyes were topically covered with Tobradex (Alcon, Fort Worth, TX, USA) ophthalmic antibiotic ointment to reduce inflammation, and 0.1 mg/kg of buprenorphine was subcutaneously injected to minimize discomfort following surgery.

### Retinal imaging

The animals were anesthetized, as described earlier in the study. Following the anesthesia induction, the pupils were dilated and cyclopleged with 2.5% phenylephrine (Akron, Inc., Lake Forest, IL, USA) and 1% tropicamide (Akron, Inc., Lake Forest, IL, USA). Local anesthetics were applied with one drop of 0.5% tetracaine (Alcon, Fort Worth, TX, USA) to lessen the blinking reflex.

#### (i) Scanning laser ophthalmoscopy (SLO)

The *en face* view of the retina was obtained using a customized scanning laser ophthalmoscope (SLO) with a 55° lens (Heidelberg Engineering, Germany) to assess the healthy retina (undamaged/pre-damaged) and damaged retina (ATP/retinal detachment), before and after transplantation. First, up to 1 month prior to inducing damage, near-infrared (NIR; 820.5 nm) reflectance images were collected across a large retinal region nasal to the optic nerve to ensure sufficient baseline coverage. The 15 damaged retinas were all imaged 1 week prior to cell transplantation with both NIR reflectance and GFP fluorescence SLO (fSLO; 488 nm excitation, 500-550 nm transmission filter). The 7 undamaged retinas were also imaged with both NIR reflectance and fSLO before cell transplantation. Post-transplantation, all eyes underwent weekly assessment using NIR reflectance and fSLO.

#### (ii) Optical coherence tomography (OCT)

Cross-sectional imaging of the retina was performed using a Bioptigen Envisu R2310 spectral domain-OCT equipped with a rabbit imaging bore (Leica Microsystems, Wetzlar, Germany). For all imaging sessions, vertical and horizontal volume scans were acquired using 650 A-scans/B-scan and 300 B-scans per volume. Line scans were acquired at the same location using 650 A-scans/B-scan with 100 repeated B-scans. For both damaged and undamaged animals, pre-transplantation baseline imaging was performed 1 week before transplant. Following cell transplantation, imaging of transplanted regions was performed weekly. Temporal changes were assessed by aligning baseline images to post-transplant images. To evaluate the overall pre-transplant retinal environment and outer retinal integrity in the two damage models relative to the undamaged model, outer retinal thickness was measured from baseline scans. In damaged retinae, thickness was extracted from retinal locations that were as close to the most damaged areas as possible while also retaining identifiable lamination. The ImageJ TurboReg plugin was used to process the line scans by registering 20 B-scans from each line scan acquisition to a selected reference frame (Thévenaz et al., 1998). The final processed image was generated via an average intensity z-projection of the registered image stack. The post-processed scans were then loaded into a custom software (Duke Optical Coherence Tomography Retinal Analysis Program; DOCTRAP) to semiautomatically segment the outer plexiform layer and the retinal pigmented epithelium (RPE) (Chiu et al., 2010, Chiu et al., 2015). The difference (in pixels) between these two layers was extracted and converted to microns for outer retinal thickness measurements. Outer retinal thickness measurements were averaged across all eyes for each of the three models: undamaged (n=7), retinal detachment (n=11), and ATP (n=4). The differences in average outer retinal thickness were compared using 2-Way ANOVA with multiple comparison tests across three models.

#### (iii) Adaptive optics scanning light ophthalmoscopy (AOSLO)

High-resolution cellular imaging was performed using a custom-built AOSLO system as previously described (Sajdak et al., 2016, Sajdak et al., 2019). Briefly, the AOSLO system uses a 785 nm superluminescent diode (SLD) (ThorLabs, Newton, NJ, USA) for NIR structural imaging and an 850 nm SLD (Superlum, Cork, Ireland) for wavefront sensing. Wavefront aberrations of the eye were measured with a Shack-Hartmann wavefront sensor and corrected with a deformable mirror (7.2 mm diameter, 97-actuator) (ALPAO; Montbonnot-Saint-Martin, France). The fSLO images from the latest time-point were utilized as a guide to locate the transplanted region in the AOSLO imaging session. Multiple overlapping images were obtained along retinal vessels and into portions of the transplanted regions. From these images, a small montage was created to facilitate alignment with the wide-field SLO and OCT imaging described above. Confocal and non-confocal AOSLO images were acquired simultaneously. AOSLO imaging was performed prior to inducing retinal damage, pre-transplant, and 5-8 weeks post-transplant, including at the endpoint of the experiment. Transplanted areas in two animals (3 eyes) were also assessed with fluorescence AOSLO (fAOSLO). Briefly, a NKT SuperK super-continuum light source equipped with a VARIA tunable filter (NKT Photonics/Hamamatsu, Bregnerødvej, Denmark) set to 488±30nm was used for GFP fluorophore excitation and imaging. The emitted GFP signal was isolated using a Semrock Brightline Fluorescence bandpass (530/55nm) filter and detected in one confocal and four non-confocal visible PMT channels. Confocal and non-confocal NIR images were also collected simultaneously during fAOSLO imaging.

### Human retinal organoid differentiation

The hiPSC-derived retinal cells used in this study were differentiated from hiPSC harboring CAG-GFP pan-expression construct at the safe harbor locus (Pei et al., 2015) at the University of California San Francisco (UCSF). All the human stem cell experiments were approved by the UCSF Institutional Review Board and Human Gamete, Embryo and Stem Cell Research (GESCR) committee. The hiPSCs cultured on a 1% Matrigel-coated plate with mTeSR plus medium (100–0274, Stemcell Technologies, Canada) were differentiated towards retinal fate *via* the embryoid body (EB) and three-step (3D-2D-3D) approach as described in the previous publication (Arthur et al., 2022, Bachu et al., 2022). hiPSCs, dissociated as small clumps, were cultured in a suspension plate with neural induction medium to form EBs. Brief exposure of EBs to varying concentrations of BMP4 (1.5-0.75-0.375 nM) in neural induction medium was carried out from Day 6-15 of retinal differentiation. BMP4-treated-EBs were then plated onto the Matrigel coated culture plate with retinal differentiation medium (RDM) for two weeks to form optic vesicles. The microscopically visible optic vesicle was manually excised and cultured in a suspension plate with 3D-RDM supplemented with 1 μM ATRA (R2625, Sigma-Aldrich, St. Louis, MO, USA) to generate 3D-retinal organoids. Medium change for retinal cultures with RDM/3D-RDM was done at every 2-day intervals until the retinal organoids were 60 (+/− 3) days old. 24 hours prior to the transplantation day, retinal organoids were treated with a small molecule inhibitor of the Notch pathway, PF-03084014 hydrobromide (PF, 10 μM) (PZ0298, Sigma-Aldrich, St. Louis, MO, USA) for 24 hours to drive the differentiation of remaining retinal progenitor cells within the organoids towards the cone fate (Figure S1) (Chew et al., 2022). Post 24-hour PF treatment, the GFP expressing cone-rich 3D retinal organoids were collected in 15 mL conical tube with fresh 3D-RDM containing ATRA. The tubes were packed with warm packs and shipped overnight from UCSF to MCW for transplantation.

### Retinal organoid dissociation for transplantation

The cone-rich 3D retinal organoids derived from human iPSC were dissociated upon receipt at MCW for transplantation. The tubes with organoids were placed in the incubator at 37°C, 5% CO_2,_ and 95% humidity by loosening the tube cap for 10 minutes immediately upon receipt of the package. Following this, the organoids were washed with 1X PBS at 50x g for 5 minutes. The supernatant was discarded, and organoids were treated with 5 mL papain solution containing Earle’s Balanced Salt Solution and 250 μL DNAs (LK003150, Worthington Biochemical Corporation, Lakewood, NJ, USA) for 30-45 minutes with gentle agitation and intermittent trituration until complete dissociation. The dissociated cell suspension was passed through a 100 μm cell strainer (22-363-549, Fisher Scientific, Pittsburg, PA, USA). The cells were enumerated using 0.4% trypan blue dye (Thermo Fisher, Waltham, MA, USA) on a hemocytometer. Finally, the cell suspension was centrifuged at 200x g, and the pellet was resuspended in the 3D-RDM with ATRA at a desired concentration of 3.5X10^4^ cells / μL for injection.

### Retinal cell transplant

The dissociated cells were placed in warm packs and taken to a dedicated surgical suite in the animal facility. A heated surgical table was covered with sterile surgical drapes and used to maintain the tree shrew’s body temperature throughout the surgery. The prepared sterile surgical equipment used was identical to that used in the experimental retinal detachment, as mentioned previously. The animals were anesthetized, and both eyes of the animals were dilated and cyclopleged as mentioned previously. Following the setup of the sterile surgical field and animal position, the intraoperative surgical scope (Proveo 8) view was optimized for both the fundus and OCT view of the target eye. The location of the damaged region was first located under the surgical scope to view the orientation and direction prior to inserting the needle. Following the preparation, 0.7-1 million dissociated cells (resuspended in 25 µL of complete 3D-RDM) were drawn up using the 33-gauge needle and 100 μL Hamilton syringe and delivered as cell bolus into the subretinal region of the undamaged and damaged tree shrew retina. Following the injection, a confirmatory OCT scan was obtained at the location of the injection to confirm the subretinal delivery of cells. Then, the animal was recovered as previously described. For all cell transplant sessions, vitrectomy was not performed.

### Enucleation and immunohistochemistry

The tree shrew eyes with successful transplants and undamaged eyes were collected for histology at the endpoint of the experiment. The tree shrews were euthanized, and the eyes were enucleated. The enucleated eyes were first rinsed in fresh and cold 1X PBS. A small cut was made along the limbus of the eye using a 20-gauge trocar to allow the fixatives to diffuse inside the eye. The whole eye globe was fixed in 4% paraformaldehyde (PFA) in 1x PBS overnight at 4°C. On the following day, the eyes were taken out of the 4% PFA solution and rinsed in 1X PBS thrice at 15-minute interval each. Next, the small patch of the transplanted retinal regions within the eye was located under the dissection scope and cut out. The small cut-out transplanted retinal patches were again immersed in 4% PFA for 20 minutes at 4°C to ensure effective fixation of the retina. The fixed retinal patches were washed with 1X PBS thrice at 5-minute intervals each. The samples were then equilibrated through 15% and 30% sucrose solution for 1 hour each or until the tissue sunk to the bottom of the tube. Finally, the sample tissue was embedded in Tissue Tek OCT compound mixed with 20% sucrose solution mixture (1:2) using a dry ice bath. 8 μm thick sections were collected on the Superfrost Plus glass slides then cryopreserved at −80°C until staining.

The frozen sections were taken out of −80°C freezer and thawed at room temperature. The sections were washed in 1X PBS for 5 minutes and permeabilized with 0.1% Triton X-100 (VWR, Radnor, PA, USA) made in 10% normal donkey serum (NDS) (Sigma Pharmaceuticals, North Liberty, IA, USA) for 15 minutes. Following permeabilization, the tissue sections were then blocked with 10% NDS (made in 1X PBS) for an hour at room temperature. Next, the primary antibodies (Table S3) diluted in 10% NDS were applied to the sections and incubated overnight at 4°C. The next day, the sections were washed in 1X PBS thrice for 5 minutes each at room temperature.

Secondary antibodies (Table S4) diluted in 10% NDS were added to the sections for 1 hour in the dark and at room temperature. After 1-hour, secondary antibodies were removed and 1 µg/mL DAPI (Roche, Basel, Switzerland) diluted in 1X PBS was applied on sections for 10 minutes in the dark at room temperature. Lastly, the sections were washed with 1X PBS thrice at 5-minute intervals each and mounted with fluoromount-G (Electron Microscopy Science, Hartfield, PA, USA) and cover glass (VWR, Radnor, PA, USA). Fluorescent microscopy images were collected using an LSM 700 inverted confocal microscope (Carl Zeiss, Oberkochen, Germany). The images were processed with ImageJ (NIH, Bethesda, MD, USA) and cropped and scaled with Photoshop CS6 (Adobe, San Jose, CA, USA).

### Cell quantification

In undamaged and detached eyes, groups of adjacent sections were stained with HuNu+ marker and with OTX2, HuC/D, or Brn3A markers. A custom software (*Mosaic*, Translation Imaging Innovations, Hickory, NC, USA) was used to manually count each cell as shown in Figure S2. For HuNu/OTX2 section, all HuNu+ cells were first counted within the subretinal space. Then, the same cell coordinates were placed onto the corresponding OTX2 image of the section to extract the colocalized cells within the subretinal space. For the inner retina and ganglion cell layer, all HuNu+ cells in the inner retina and ganglion cell layer were first counted. Then, the number of cells that colocalized with HuC/D or Brn3A marker was extracted. The percentage of OTX2, HuC/D, or Brn3A makers that colocalized with HuNu+ cells was computed (Graphpad Prism, San Diego, CA, USA).

## Results

### ATP and retinal detachment induce outer retinal damage

To assess how transplanted hiPSC-derived retinal cells would respond in a damaged environment, we first sought to induce outer retina-specific degeneration via chemical and physical methods. Intravitreal injection of ATP has proven to be effective for inducing photoreceptor-specific degeneration in numerous animal models, including mice, cats, and 13-LGS,(Puthussery and Fletcher, 2009, Aplin et al., 2014, Follett et al., 2024) where ATP induces the death and loss of photoreceptors through excitotoxicity by the overactivation of P2X7 channels (Puthussery and Fletcher, 2009). The alternative method of inducing photoreceptor degeneration was done through experimental retinal detachment by subretinal injection with normal saline. This method physically separates photoreceptor and RPE from each other, leading to photoreceptor degeneration. Additionally, a previous study has shown that normal saline can induce damage to cells within the outer retina, including photoreceptors and RPE (Hombrebueno et al., 2014). We first evaluated the detachment model in our earlier trials (Table S1, Eye 1-4). Following the assessment of the detachment model and transplant, we then also assess the ATP model in the later batch with the detachment model (Table S1, Eyes 16, 18, 20, 22). In total, 4 eyes had ATP-induced damage, and 11 eyes had retinal detachment among the 12 animals used in the study, with three control animals being kept undamaged (ATP, n=4; Detachment, n=11; Undamaged, n=7 eyes).

Of the four eyes that received intravitreal ATP injections, two eyes that were injected with an effective dose of either 0.01 M or 0.0085 M ATP, developed retinal damage within 1 week post-injection. The other two eyes that received an effective dose of 0.0075 M ATP failed to develop degeneration. These two eyes were re-injected with a higher dose of 0.0085 M ATP, which successfully induced photoreceptor damage by 1 week post-injection. For the experimental retinal detachment model, the successful detachment of the photoreceptors from the RPE was documented at the time of injection through intraoperative OCT. Photoreceptor degeneration was observed within 1 week post-injection in both the ATP and retinal detachment models using multimodal *in vivo* imaging (Figure 1). All damaged injection parameters are listed in Table S1. The quantification of outer retinal thickness was measured between the undamaged, retinal detached, and ATP-induced models using OCT scans collected 1 week prior to transplantation (Figure S3, S4). The outer retina was significantly thinner in the ATP-damaged retinas compared to the undamaged controls (p=0.01, Tukey’s multiple comparison test), though the experimentally detached retinas did not have significantly different outer retinal thickness.

**Figure 1.**
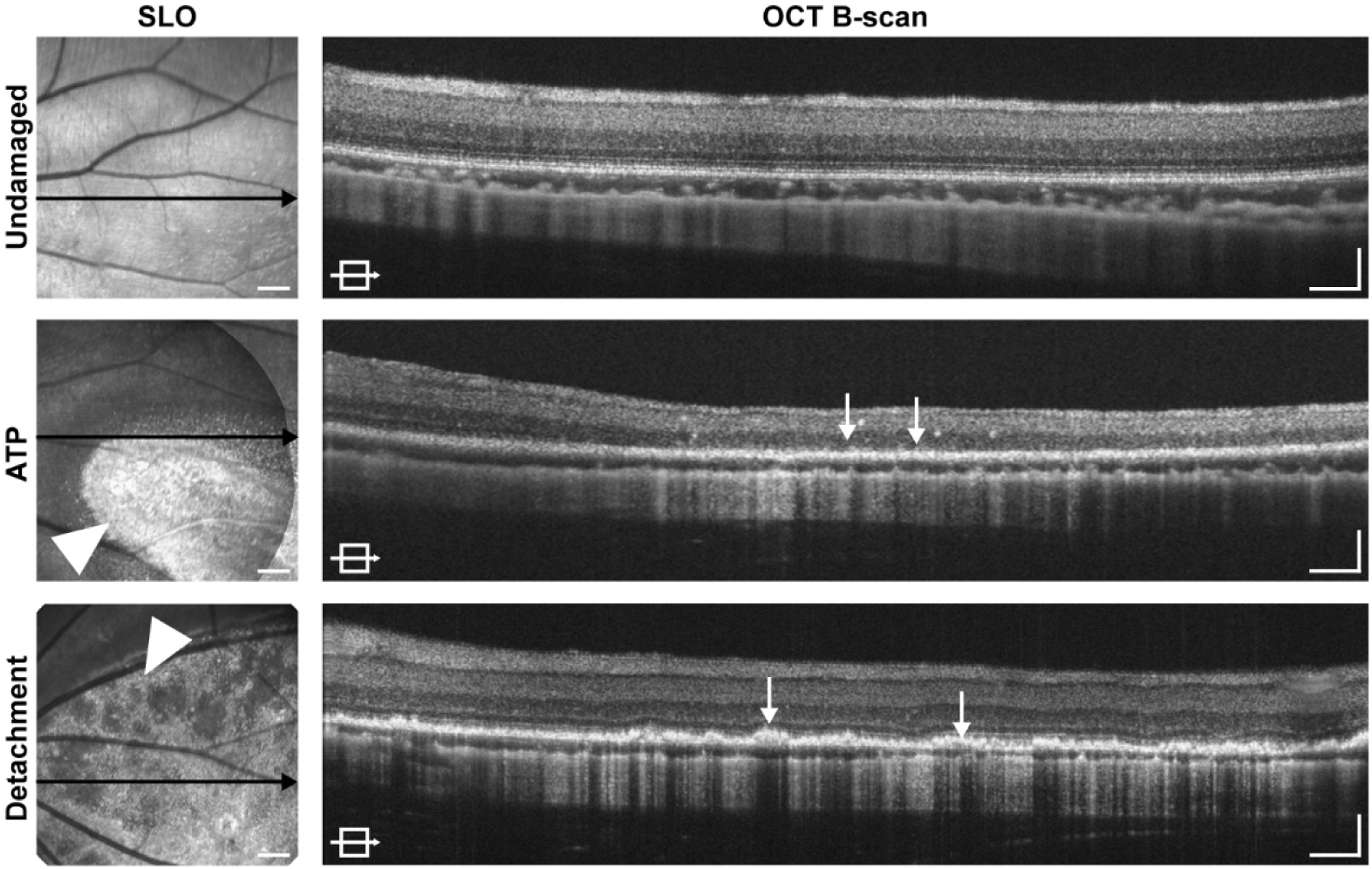
Retinal degeneration induced via ATP or retinal detachment in comparison to undamaged tree shrew retinae. Representative SLO fundus and cross-sectional OCT B-scan images of the undamaged (7A4D956_OD) and damaged regions in ATP (7E9FAAD_OD) and detachment model (7D642BC_OD) of tree shrew retinae 1 week prior to transplantation. Both damage models show altered reflectivity on the SLO near-infrared reflectance (NIR) images (arrowheads) and disrupted retinal lamination (arrows) on the OCT images. Scale bars are 5 degrees (SLO image), 100μm (OCT B-scan, axial), and 2 degrees (OCT B-scan, lateral).

### Transplanted cell survival is poor under oral cyclosporin A immunosuppression

We initially adapted the immunosuppression protocol used in our previous 13-LGS study (Yu et al., 2024). Following the initiation of immunosuppression, GFP-expressing hiPSC-derived retinal cells were delivered to the subretinal space of 10 eyes. We transplanted undamaged (n=5) eyes and eyes damaged via retinal detachment (n=5). Follow-up fSLO demonstrated the presence of GFP signal at transplanted regions, and the OCT B-scan confirmed that the cells were delivered subretinally. However, the transplanted regions lost all GFP expression by four weeks (Figure S5). We also observed retinal layers began developing additional lesions phenotypically-similar to retinal delamination (Figure S5). While we cannot rule out that this can be of mechanical origin, it is likely due to rejection of cell transplantation as this was not observed in any the injected eyes following modification to immunosuppression protocol.

### Transplanted cells localize subretinally and survive for up to eight weeks in tree shrew retinae under species-specific immunosuppression protocol

We next aimed to apply a species-specific immunosuppression protocol in order to minimize cell rejection, as has been shown for canine transplants (Ripolles-Garcia et al., 2022). With the multi-drug immunosuppression protocol containing cyclosporin A, methylprednisolone, doxycycline, sirolimus, azathioprine, and basiliximab, we transplanted retinal cells in 12 additional eyes. Following the induction of photoreceptor degeneration, we transplanted GFP-expressing hiPSC-derived retinal cells subretinally into 12 eyes: undamaged (n=2), ATP-damaged (n=4), and detachment-damaged (n=6). Amongst the 12 eyes injected with hiPSC-derived retinal cells, three eyes (ATP, n=1; retinal detachment, n=2) had failed injections, where cells were delivered into the vitreous. Through fSLO imaging, we tracked the cell survival by observing the GFP expression of the transplanted hiPSC-derived retinal cells. One animal failed to recover from anesthesia during a follow-up imaging session. In this case, the two eyes (ATP, n=1; retinal detachment, n=1) were collected at four weeks post-transplantation and processed for immunohistochemical analysis. In transplants in undamaged eyes, we observed persistent GFP expression by fSLO up to 5 weeks post-transplantation (Figure 2A). Under damaged condition models, to assess the durability of transplanted cells, we extended our follow-up imaging time points for up to 8 weeks (Figure 2A). For the damaged models (both ATP and retinal detachment), we observed GFP-expressing cell survival for up to 8 weeks (Figure 2A). Additionally, the cross-sectional OCT images corresponding to the GFP signal observed on fSLO confirmed the sub-retinal localization of the transplanted cells in these regions in all nine eyes (ATP, n=3; retinal detachment, n=4; undamaged, n=2) (Figure 2B).

**Figure 2.**
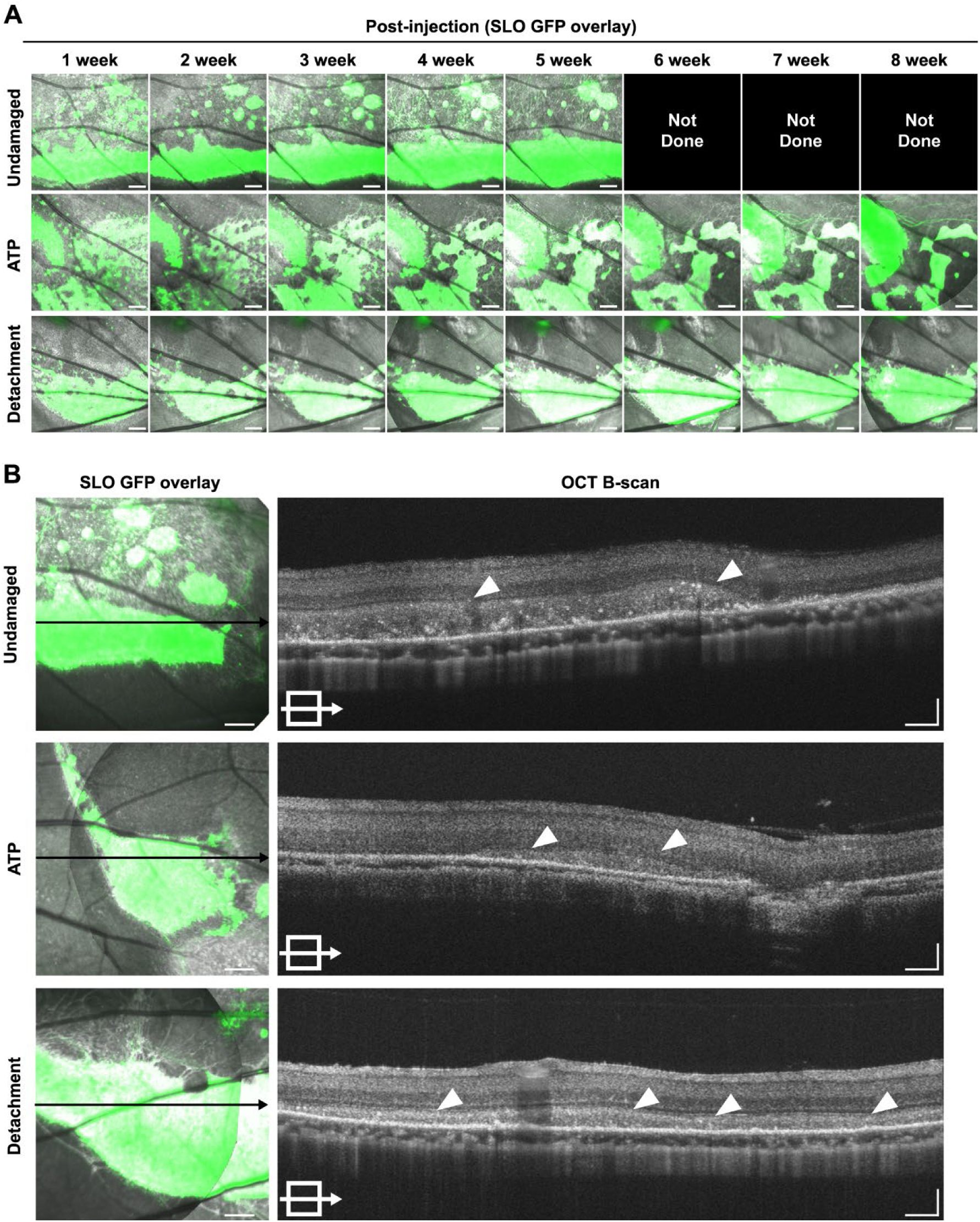
Transplanted cell survival and localization by *in vivo* imaging modalities. (A) Representative SLO images of undamaged (7A4D956_OD), ATP (80F0D53_OD), and detachment (7E6CA55_OS) models of tree shrew retina showing the GFP overlay on the near-infrared SLO image from 1-week until 8-weeks of cell transplantation. (B) Representative OCT B-scans (right) through the locations of GFP signal (cell transplanted, black horizontal line) on the SLO (left) of undamaged, ATP, and retinal detached animals. White arrowheads in OCT scans indicate subretinally localized transplanted cells. Scale bars are 5 degrees (SLO images), 100μm (OCT B-scan, axial), and 2 degrees (OCT B-scan, lateral).

### AOSLO imaging reveals presence of photoreceptor-like structures in transplanted regions

We further characterized the transplanted region *in vivo* using single-cell resolution adaptive optics SLO (AOSLO) imaging. Photoreceptor signals detected using confocal AOSLO imaging are thought to represent intact outer segment waveguiding, while photoreceptor signals on the non-confocal modality are thought to originate from multiply-scattered light from the photoreceptor inner segments (Scoles et al., 2014). Using the widefield fSLO images of transplanted GFP-expressing hiPSC-derived retinal cell locations for navigation, confocal and non-confocal AOSLO videos were collected to create a contiguous montage spanning an area from within the transplant to an area outside of the transplanted region in each eye. Representative AOSLO montages from all three models (undamaged, ATP, and retinal detachment) are overlayed on top of their respective fSLO images (Figure 3A, 3D, 3G). In undamaged retinas, structures resembling the host cone photoreceptor outer segments were observed in confocal images collected outside the transplanted region (Figure 3B). No structures resembling outer segments were found in confocal images from regions within transplanted cell patches in any of the three models (Figure 3B, 3E, 3H). This could be due to the stage of maturation of hiPSC-derived retinal cells (∼100 days, 8 weeks post-D60 organoid-derived transplants) when they are not expected to develop outer segments (Capowski et al., 2019). Similarly, structures resembling host inner-segments were observed in non-confocal images from areas outside transplanted regions in all three models (Figure 3C, 3F, 3I). Non-confocal images showed additional structures that were less photoreceptor-like in appearance, both within and outside of transplanted areas.

**Figure 3.**
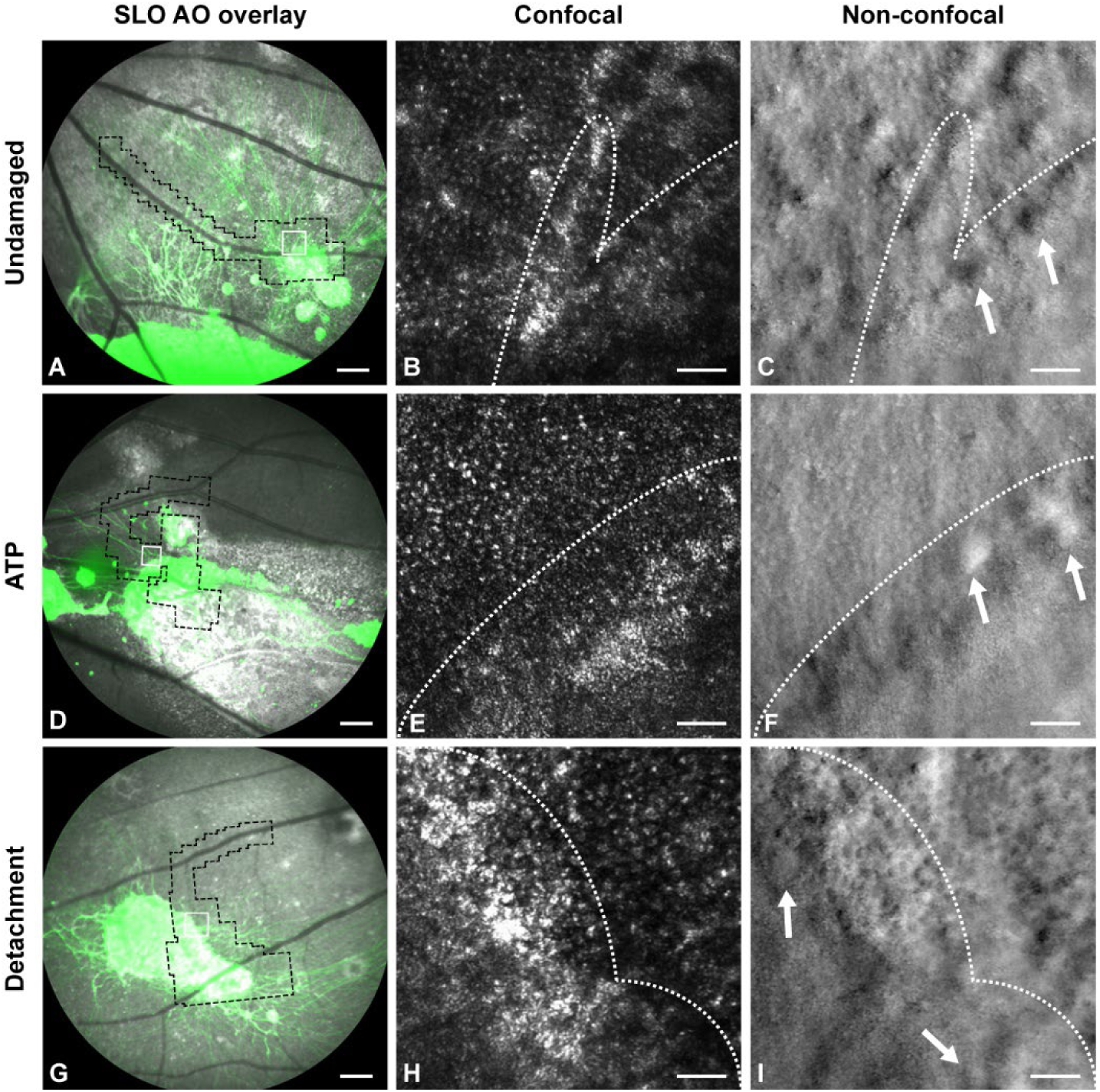
AOSLO imaging of undamaged and damaged tree shrew retinae. (A, D, G) Representative SLO GFP overlay images showing the location of the AOSLO montages (black dotted outline) for undamaged and damage models (ATP and detachment). (B, E, H) Representative AOSLO images in the selected region of interest (white box within panels A, D, G) at the border of the transplanted graft for the undamaged, ATP, and detachment models in both confocal (B, E, H) and non-confocal (C, F, I) modalities. Scale bars are 5 degrees (SLO fundus images) and 30 μm (AOSLO images).

Structures within transplanted areas were further evaluated in two animals (3 eyes) using confocal and non-confocal fluorescence AOSLO (fAOSLO), which was designed to capture GFP fluorescence signal. We confirmed that GFP signal co-localized between fSLO and fAOSLO images, including areas of speculated neurite sprouting as well as larger, diffuse regions of GFP signal. (Figure S6). The fAOSLO, which only collects GFP signal, showed photoreceptor-like structures similar to those in the NIR AOSLO images. In one location in the ATP model, the non-confocal fAOSLO GFP signal contained a patch of contiguous structures of similar size and organization to the remnant host inner segments seen in the NIR non-confocal images outside the transplanted region (Figure S6). Strong but diffuse fAOSLO signal was found in other transplanted locations, both with and without photoreceptor-like structures also being visible. From the fluorescent signal that was observed in the fAOSLO non-confocal channels, the observed GFP signal could either be from transplanted cells with developed inner segments or there could have been remnant host cells Secondly, it could also be that the cells were transplanted underneath the patch with remnant host cells, and the cellular structure belongs to the host cells above the GFP expression transplants.

### Transplanted cells express human-specific and retinal-specific markers

At eight weeks post-transplantation, the animals were euthanized for histological analysis. The retinal sections were processed and stained with anti-GFP (due to tissue fixatives quenching the transplant’s GFP during processing) and DAPI to locate the transplanted graft in undamaged (Figure 4A, 5A, 5E), retinal detachment (Figure 4E, 5I, 5M), and ATP-treated (Figure S7) eyes. Next, we sought to confirm the human origin of the transplanted cells by staining for expression of human-specific markers, HuNu, and LMNB2 (ATP) (Yu et al., 2024). A majority of HuNu+ cells that colocalized with GFP+ cells were in the subretinal space (SRS) of undamaged (Figure 4A, 4B, 4D, 5A, 5B, 5D), detached (Figure 4E, 4F, 4H, 5I, 5J, 5L), and ATP-treated (Figure S7) eyes. As described in our prior study in 13-LGS, some human marker-expressing cells lose their GFP expression.

**Figure 4.**
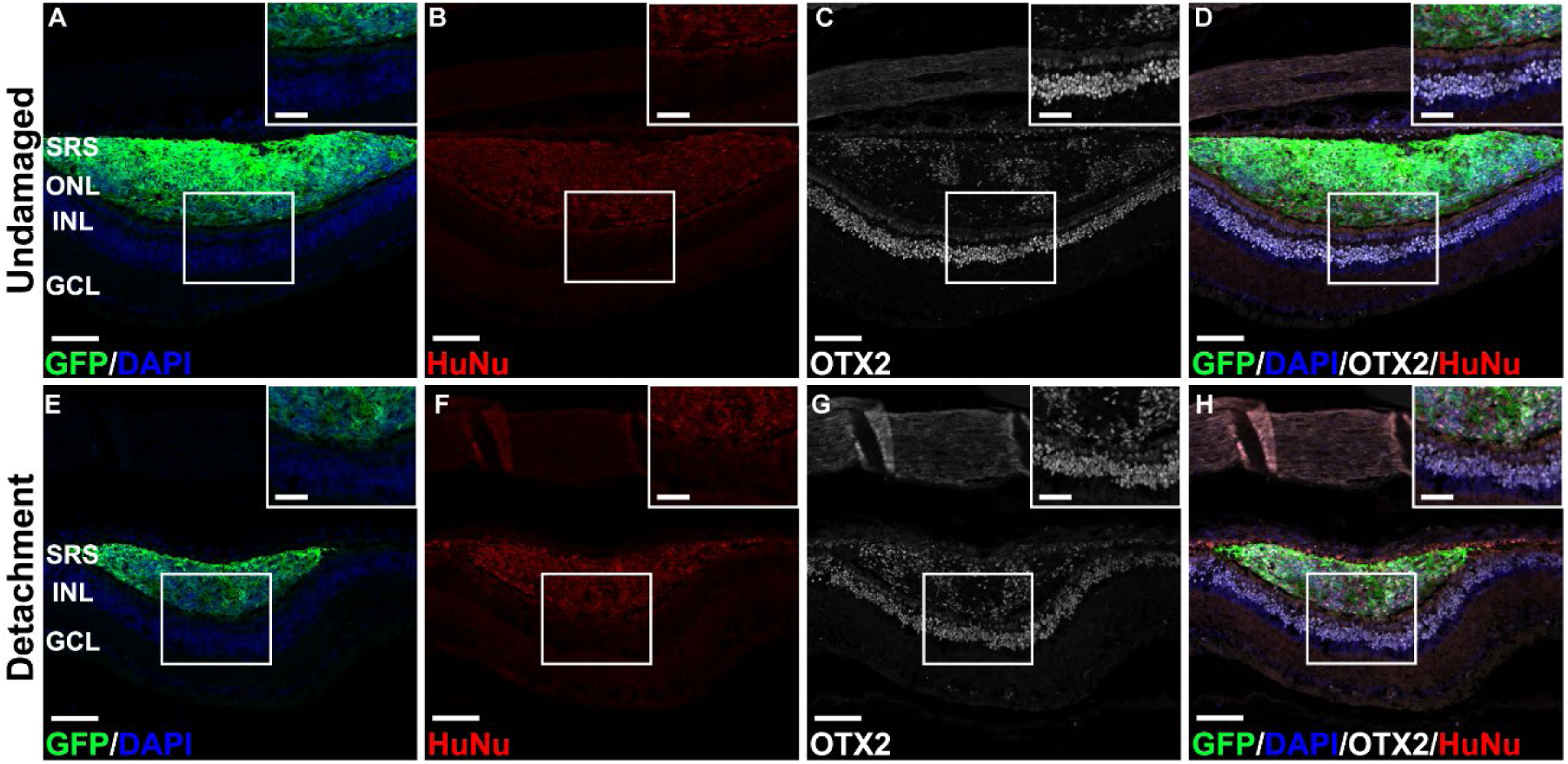
Histological analysis confirms the human origin and photoreceptor fate of the transplanted graft in the undamaged and retinal detached retina. (A-H) Representative immunofluorescence images of the undamaged and retinal detached eyes with transplanted grafts showing the location of the GFP+ cells (A, E), HuNu expression at the transplant location (B, F), OTX2 expression across the retina (C, G), merge of GFP, DAPI, OTX2, and HuNu (D, H). Scale bars are 100 μm (large image) and 50 μm (inset image). SRS, subretinal space; ONL, outer nuclear layer; INL, inner nuclear layer; GCL, ganglion cell layer.

To assess the fate of the transplanted cells, we stained retinal sections with a photoreceptor and bipolar marker (OTX2) (or CRX for ATP). The majority of cells within the SRS were OTX2+ and colocalized with both GFP and HuNu markers in undamaged (Figure 4C, 4D), detachment model (Figure 4G, 4H), and ATP-damaged (Figure S7) eyes. Marker colocalization was quantified as the percent of HuNu+ cells that were also OTX2 positive in the subretinal space of undamaged (51.9 +/− 3%) and retinal detached models (60.4%). (Table S2). Quantification of the ATP-damaged eyes was not possible due to severe degeneration that prevented sufficient processing for additional markers.

Several HuNu+ cells were observed in the inner retina in transplanted, undamaged, and detached eyes. The retinal sections of transplanted regions were labeled with HuC/D and Brn3A markers, which label ganglion and amacrine cells. The majority of the inner retina HuNu+ cells colocalized with HuC/D in the INL and ganglion cell layer (GCL) of undamaged (85%) and retinal detachment (100%) models (Figure 5C, 5K and Table S2). These infrequently expressed ganglion cell markers, Brn3A (46.9%) in undamaged and (33.3%) in the retinal detachment model, suggest that the majority of HuNu+ cells that migrate into the inner retina are likely amacrine cells (Figure 5F, 5G, 5N, 5O and Table S2).

**Figure 5.**
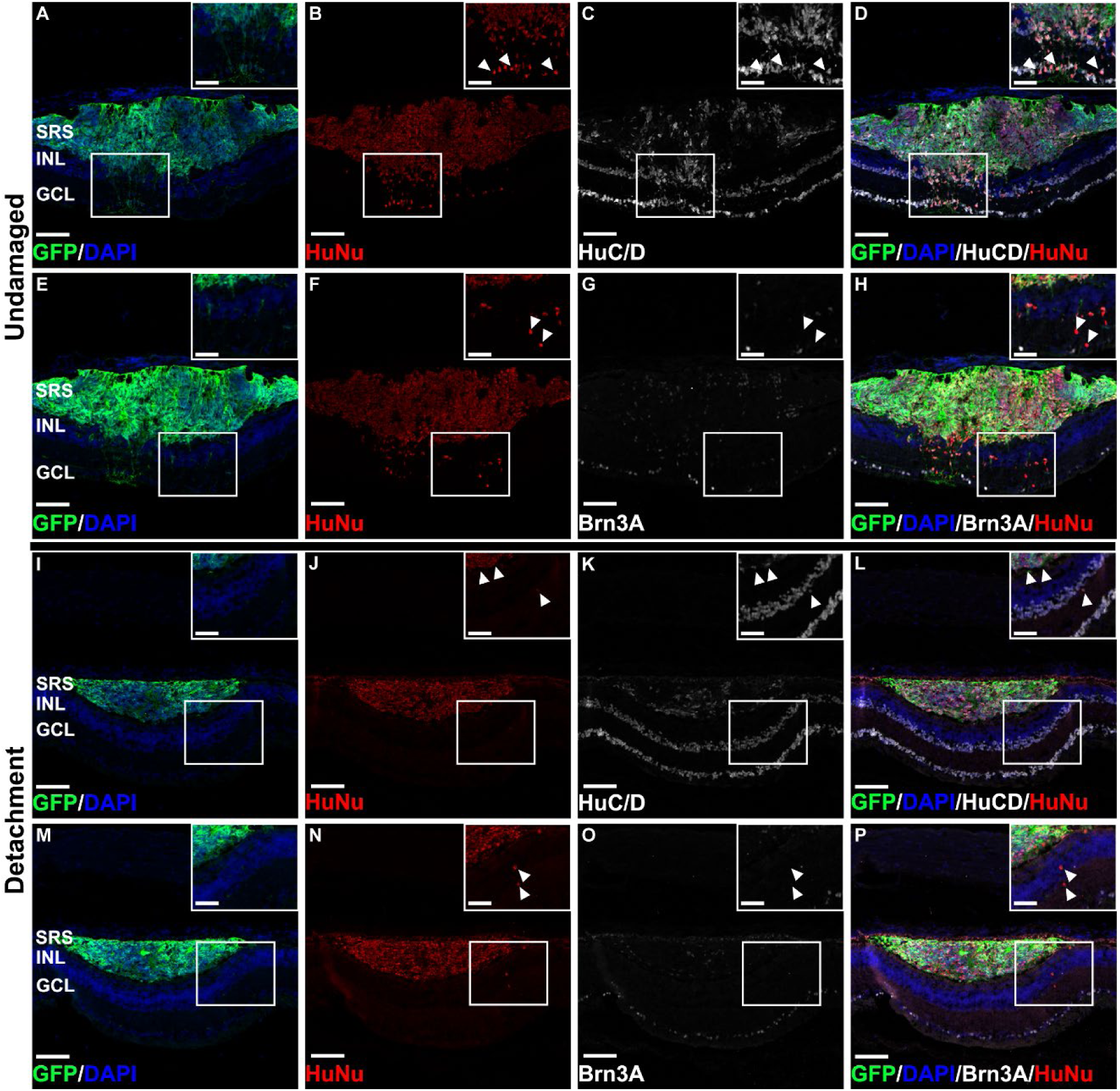
Histological analysis confirms the retinal ganglion or amacrine cell fate of transplanted human cells in the inner retina of the undamaged and detached tree shrew eyes. Representative immunofluorescence images of transplanted undamaged and detached retina showing the location of the GFP+ transplanted graft (A, E, I, M), HuNu+ human origin cells (B, F, J, N), HuC/D + retinal ganglion and amacrine cells (C, K), Brn3A+ specific retinal ganglion cells (G, O) and merge of each marker (D, H, L, P). Scale bars are 100 μm (large image) and 50 μm (inset image). SRS, subretinal space; INL, inner nuclear layer; GCL, ganglion cell layer.

## Discussion

There are a number of ongoing efforts toward optimizing cell therapy to treat end-stage retinal degeneration (MacLaren et al., 2006, Aboualizadeh et al., 2020, Ripolles-Garcia et al., 2022, Zhu et al., 2017). Numerous studies describe variable results based on the choices of animal models and protocols, including our group’s prior transplant study in the 13-LGS, a cone-dominant animal model for assessing therapies in a cone-enriched microenvironment like that of the human fovea, (Yu et al., 2024). However, the physiology of the 13-LGS is unique as they are obligate seasonal hibernators. While our study attempted to avoid seasonal confounds by completing the work during the active season without crossing into the hibernation season, the animals’ retinae begin to undergo structural and functional changes leading up to the hibernation season (Sajdak et al., 2018). Therefore, our current study explored the survival of cone-rich hiPSC-derived retinal cells in the non-hibernating, cone-dominant tree shrew retina under both undamaged and damaged conditions. Prior to transplants in damage models, we induced retinal damage through physical and chemical methods, then documenting the extent of the damage to serve as a baseline reference. Following the transplant, we demonstrated survival of transplanted human retinal cells in all three models (undamaged, physical damage, and chemical damage) using *in vivo* clinical imaging tools, including SLO to assess cell survival and OCT to track cell localization. Furthermore, high-resolution AOSLO imaging was used to evaluate cellular structure before and after transplantation. Although AOSLO imaging is not widespread in studies like ours, there have studies that have assessed the survival of transplant cells through fluorescence and/or non-confocal AOSLO imaging (Aboualizadeh et al., 2020, Yu et al., 2024). Our histology findings also supported and validated the *in vivo* findings at the cellular level by confirming photoreceptor and bipolar cell fate with human origin markers.

Utilizing non-invasive imaging to track the longitudinal survival and localization of cell transplants increases the translatability of this study’s results to observations in human clinical trials seeking to evaluate cell-based therapies, especially those focused on rescuing cone photoreceptors (given the cone-dominant nature of our animal host). Non-invasive clinical imaging tools, such as SLO and OCT, are commonly used to monitor disease progression clinically and define clinical trial outcomes (Schwartz et al., 2015, Liu et al., 2017, Song et al., 2015). Furthermore, the high-resolution AOSLO imaging used in this study also enabled the detection of early structural changes on the cellular level, information undetectable with current clinical devices. Adaptive optics techniques are increasingly used to assess subtle, clinically undetectable changes in retinal structure in a number of human retinal diseases (Godara et al., 2010, Scoles et al., 2014, Syed et al., 2013, Langlo et al., 2013). Tools enabling early access to structural changes are critical for therapeutic development and could provide guidance as to when and where to probe for functional recovery. In this study, we characterized transplanted cell morphology and documented structures resembling cone inner segments. Future, longer-term studies will be essential to evaluate inner and outer segment development as well as probe the function of transplanted photoreceptors in these retinal regions. Functional assessments of transplanted cells may benefit from the emerging method of optoretinography, which enables measuring the function of individual photoreceptor cells (Li et al., 2025).

As previously stated, variable results could be the result of using different species, different immune-suppression protocols, and many other varying experimental conditions. One of the surprising findings was the degree of immunosuppression required to prevent cell rejection. While studies in mice, rats, and 13-LGS have shown effective survival of transplanted cells with minimal immunosuppression, (Lamba et al., 2009, Mandai et al., 2017, McLelland et al., 2018, Yu et al., 2024) we and others have shown that the immune privilege is incomplete and may be species-specific (Zhu et al., 2017, Ripolles-Garcia et al., 2022). The immune-privileged status of the retina is further affected by damage and disease, which likely drives inflammatory activation of microglia to further promote inflammatory infiltration and, thereby, a rejection response (Patel and Lamba, 2023). The integration reported in this current study in tree shrews was generally similar to the findings of our prior study in 13-LGS (Yu et al., 2024). However, we did switch to a tree shrew-specific immunosuppression protocol in order to achieve similar outcomes as 13-LGS. This further highlights our earlier point that the retina’s immune-privileged status may play an important role in the study of species immune-suppression protocols and what needs to be suppressed in addition. While we did not explore the mechanism, which is a limitation of our current study, the working conditions established in this study will provide a solid foundation for exploring changes in mechanistic detail at the molecular level of local immune microenvironmental differences.

There were minimal to no differences in transplantation conditions besides using two different cone-dominant models and their specific immunosuppression protocols from our studies. While comparing the *in vivo* findings, we did not observe epiretinal localization of cells in any of the transplanted eyes across the undamaged and damaged cohorts in our present study. In contrast, our previous 13-LGS study showed up to 17% of transplants had epiretinal localization of cells. Similar to the *in vivo* imaging outcomes, we also observed the migration of cells in the damage model of 13-LGS, which was not observed in tree shrew transplants. There are several reasons for this, such as the severity of the outer retinal damage, glial responses following degeneration, and outer limiting membrane degeneration. Data from some rodent studies have suggested that glial scarring and loss of outer limiting membrane integrity negatively impact the success rate of transplantation (Barber et al., 2012, Pearson et al., 2010). Glial scarring has also been observed to hinder the integration of cells post-transplantation in a canine model (Ripolles-Garcia et al., 2022). In a ground squirrel study, the glial response was found to be suppressed following retinal detachment (Lewis et al., 2002). While this study was done in the California ground squirrel, another study using the 13-LGS also reported possible suppression of the glial response (Merriman et al., 2016, Lewis et al., 2005). Alternatively, a glial response was observed in a retinal degeneration model in the northern TS, suggesting they do not have a suppression of glial response as seen in ground squirrels (Hu et al., 2022). The current literature suggests that the presence or absence of a glial response could be the reason why we observed the migration of cells in 13-LGS but not in TS. The difference in glia response, glia scarring, and outer limiting membrane integrity has been shown to be related to the integration and or migration of cells following cell transplant in rodents and canine models (Ripolles-Garcia et al., 2022, Barber et al., 2012). In the 13-LGS damage model post-transplant, we have observed migration of human cells into the inner retinal layers in the ATP model and, to a lesser degree, in the detachment model. However, this was not observed in either of the TS damage models. One of the possible reasons for the lack of migration of cells in the ATP model was the incomplete clearing of the outer limiting membrane and outer nuclear layer in our TS study. The ATP model was studied and optimized for 13-LGS in a separate study from our group, which optimized the induced damage model based on their dosage to more accurately and completely clear the outer nuclear layer and likely also remove the external limiting membrane in the 13-LGS model (Follett et al., 2024). Based on the 13-LGS finding, the ATP model would likely still be promising to induce outer retinal focused damage; further testing and optimization would need to be pursued in the TS model.

Further, one of the commonly discussed differences in transplantation and cell therapy studies is whether the transplanted cells can transfer their subcellular components to the host retina. This has been observed across various published rodent studies (Pearson et al., 2016, Santos-Ferreira et al., 2016, Ortin-Martinez et al., 2017, Singh et al., 2016, Decembrini et al., 2017). We did not observe any GFP+ cells outside of the delivered transplants, and the GFP+ marker colocalized with human nuclear markers. These strongly suggest a lack of material transfer between the transplanted cells and the host retina in our present study. We observed the same in our previous study using the 13-LGS model, where we also employed and validated a xenogeneic transplant strategy. A recent study reported material transfer activities between the human cells under *in vitro* culture, but the same was not observed when the human cells were co-cultured with mouse cells or when the human cells were transplanted into mice (Ho et al., 2024). Based on this study, the evidence suggests that cells are more likely to undergo material transfer in allogeneic studies. Therefore, future studies should be pursued to further trace the material transfer activity under allogenic transplant conditions in both 13-LGS and tree shrews. This would offer further insight into our understanding of material transfer and, more specifically, the possibility of material transfer from the transplanted retinal cells in cone-dominant host models.

Overall, the present study explored and assessed hiPSC-derived retinal cell survival in cone-dominant northern tree shrews. By comparing findings from two of our studies, we gained insights into possible differences in cellular responses between the two xenogeneic transplant studies. While stable cell survival was observed using the simple immunosuppression protocol in 13-LGS, the same protocol had led to cell rejection in tree shrews. The current study demonstrated that the immunosuppression protocol had to be changed to maintain the same level of cell survival in the tree shrew as in the 13-LGS. As we were able to establish stable transplant cell survival in tree shrews, the next step would be to conduct a long-term survival study in tree shrews to explore cell maturation and integration. In 3D-retinal organoid studies, photoreceptors potentially developed the functional component of the cell’s outer segment much later (∼>150 days) during the retinal differentiation timeline (Capowski et al., 2019, Kandoi and Lamba, 2023). Hence, future post-transplantation studies in tree shrews require an extended timeline to assess the long-term survival at a minimum of 17 weeks or 120 days post-transplantation.

## Author contributions

CTY, SK, JC, DAL contribute to the development, design, execution and analysis of the project. CTY, HMF, PS contributed to the *in vivo* imaging of the experiment. SK conduced the retinal organoids experiments. SK and EM conducted the immunohistochemistry experiments. RP and DML conducted the dissociation of organoids and cell viability assessments. KA and SF provided the animal health monitoring and husbandry advice throughout the project. CTY, JC, and DAL wrote the manuscript. All authors provided feedback in reviewing and editing of the manuscript.

## Acknowledgments

The authors would like to acknowledge Drs. Brian Samuels and Timothy Blekinsop for sharing the tree shrew immunosuppression protocol; Dr. Alfredo Dubra and his team for building the custom AOSLO system, continuing support on the development of image acquisition software, the AOSLO control software, and the image registration software; Dr. Jeremy Rogers and Brian Higgins for their support in the maintenance of the AOSLO system; David Schwabe and Kenneth Allen for developing the anesthesia protocol; Christine M. Skumatz, Lisa King, and the MCW biomedical research center animal facility staff for the animal husbandry; Amanda Brantner for their support with administrating the immunosuppression treatments and continuously maintain and monitor animal health; and Cassandra Martinez for sectioning tree shrew eyes.

