## Supplementary material for "Transplantation of human iPSC-derived retinal cells in the cone-dominant Northern tree shrew": Figure S1

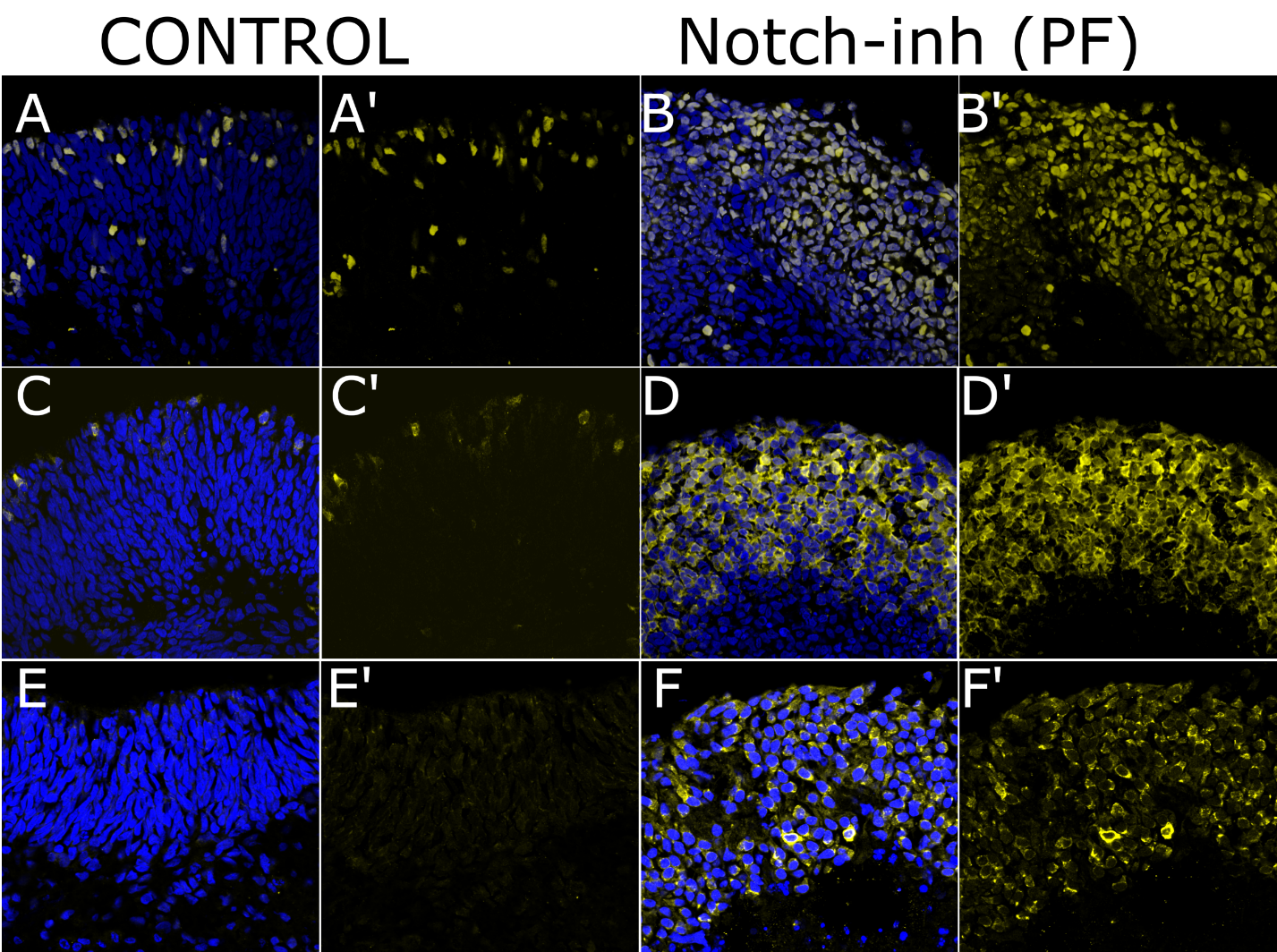

**Supplementary Figure S1. Photoreceptor differentiation in Day 60 retinal organoids following treatment of Notch pathway inhibitor.** There is significant protein expression of Crx (A, A' (control) and B, B' (Notch-i), Cone arrestin (ARR3, C, C' vs D, D') 2 weeks following treatment.
