## Supplementary material for "Transplantation of human iPSC-derived retinal cells in the cone-dominant Northern tree shrew": Figure S2

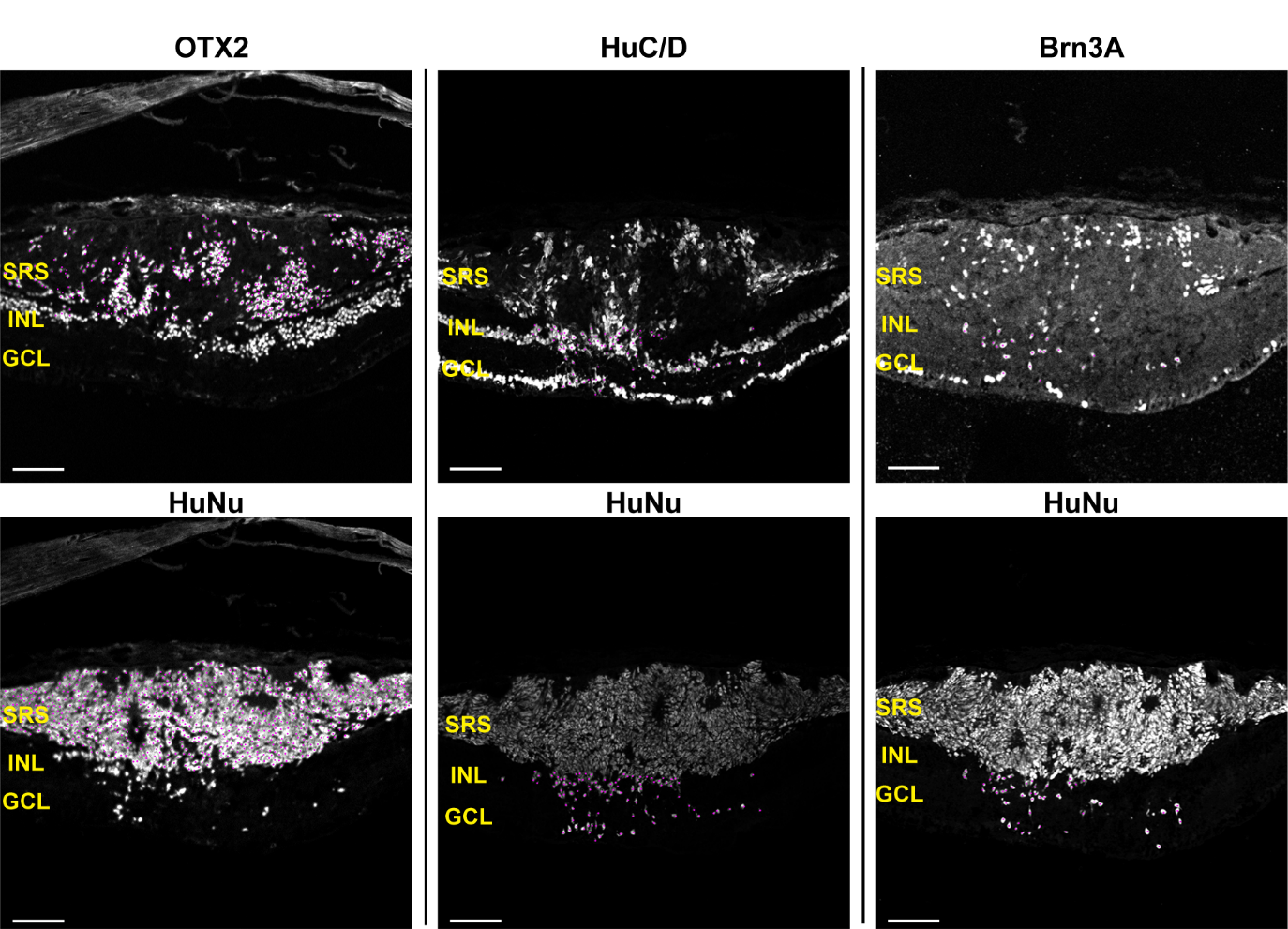

**Supplementary Figure S2. Quantitation of cell marker colocalization.** Example images of quantified sections in undamaged retina labeled with OTX2, HuC/D, and Brn3A (top row) and their corresponding HuNu images (bottom row). Scale bars are 100 μm. SRS, subretinal space; INL, inner nuclear layer; GCL, ganglion cell layer.
