## Supplementary material for "Transplantation of human iPSC-derived retinal cells in the cone-dominant Northern tree shrew": Figure S3

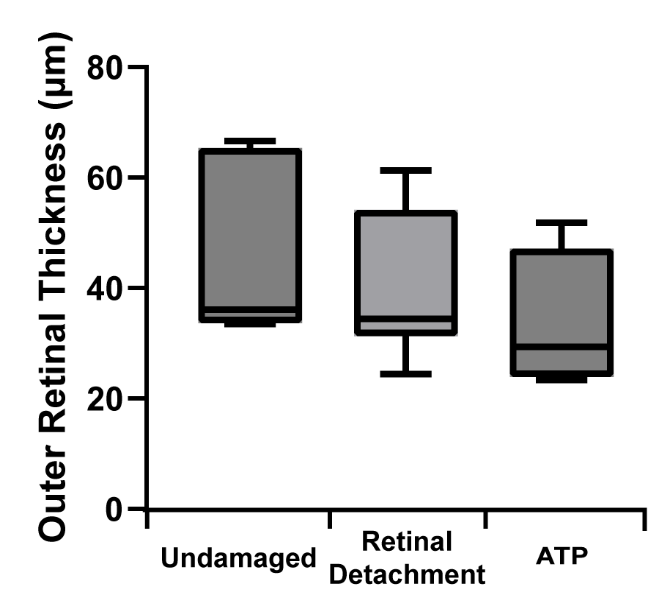

**Supplementary Figure S3. Quantification of outer retinal thickness across models.** Shown is a comparison between the undamaged (n=7), retinal detachment (n=11), and ATP models (n=4) at 1 week prior to transplantation. A significant reduction in thickness was seen in the ATP eyes compared to the undamaged eyes (Tukey’s multiple comparison test, p=0.01). The difference between the undamaged and the retinal detachment groups was not significant (Tukey’s multiple comparison test, p=0.91).
