## Supplementary material for "Transplantation of human iPSC-derived retinal cells in the cone-dominant Northern tree shrew": Figure S4

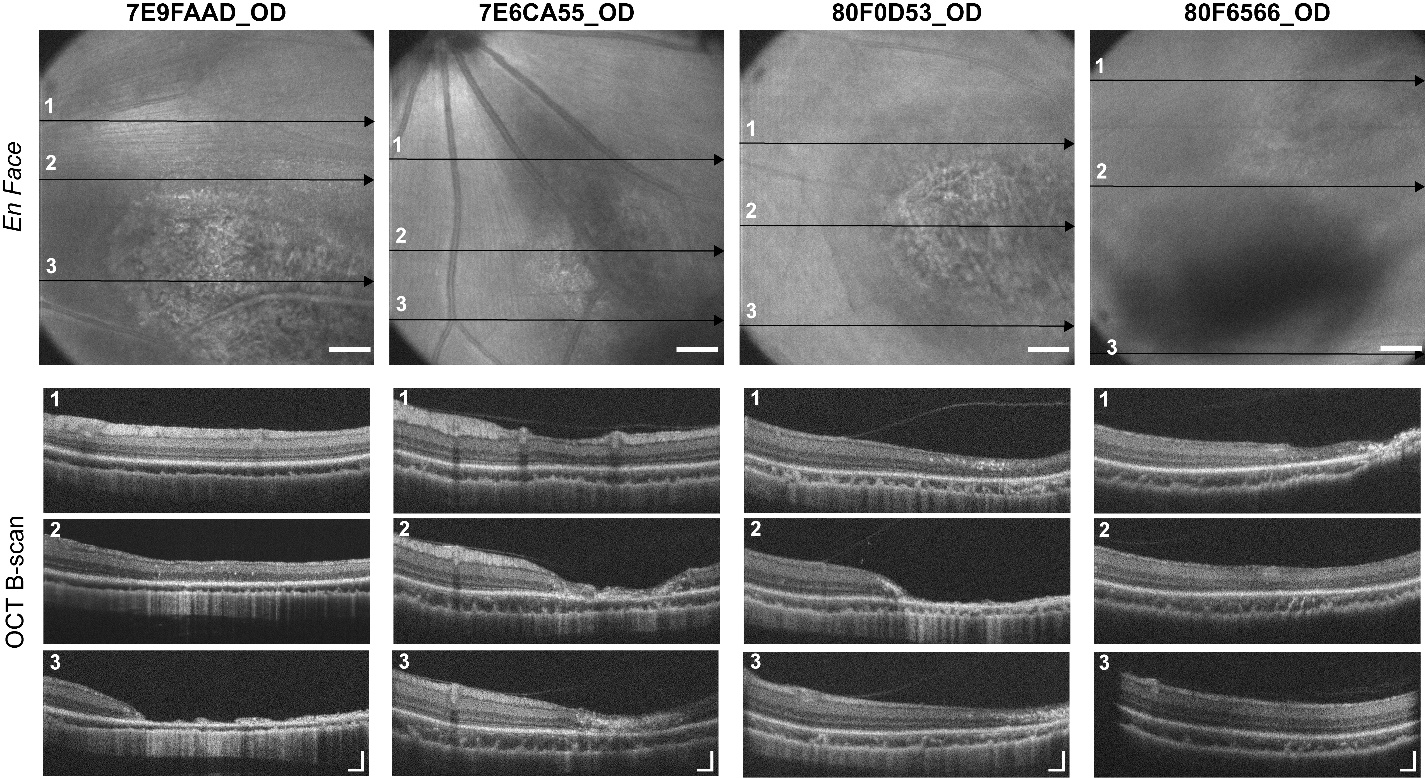

**Supplementary Figure S4. Intra-ocular variation across the ATP-induced damaged retinas 1 week before transplantation.** Representative *En Face* and OCT B-scan images of the 4 ATP-induced damaged retinas. The *En Face* image illustrates the location and extent of retinal damage. Intra-ocular variations across the retinal damage area can be seen in the OCT B-scans. The arrows indicate the specific retinal section where the respective OCT B-scan was acquired. Scale bars are 5 degrees (*En Face* image), 100μm (OCT B-scan, axial), and 2 degrees (OCT B-scan, lateral).
