## Supplementary material for "Transplantation of human iPSC-derived retinal cells in the cone-dominant Northern tree shrew": Figure S5

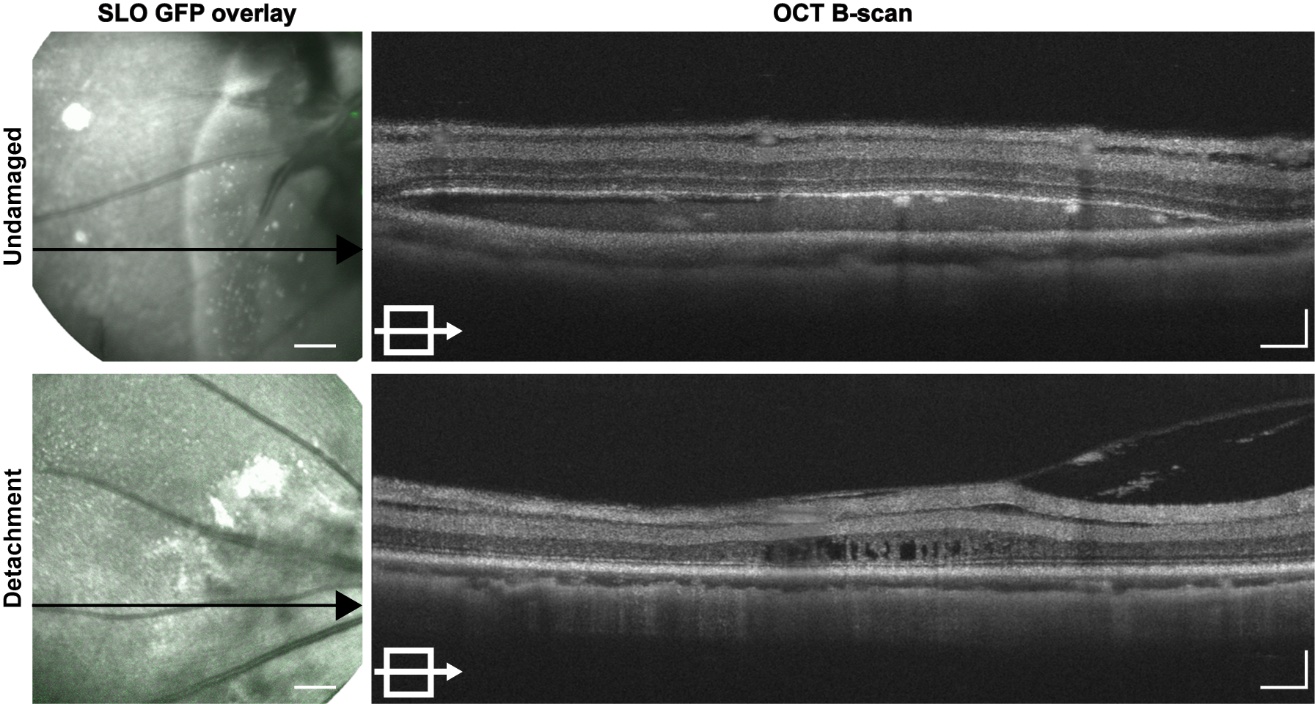
**Supplementary Figure S5. hiPSC transplantation in tree shrews treated with adapted 13-LGS immunosuppression protocol**. Representative images of the SLO GFP overlay fundus image and cross-sectional OCT B-scan from two transplanted retinae. Transplants in undamaged tree shrew retina had lost all GFP signal at 4 weeks post-transplant (top), and retina delamination was observed at 2 weeks post-transplant in the retinal detachment model (bottom). Scale bars are 5 degrees (SLO images), 100μm (OCT B-scan, axial), and 2 degrees (OCT B-scan, lateral).
