## Supplementary material for "Transplantation of human iPSC-derived retinal cells in the cone-dominant Northern tree shrew": Figure S6

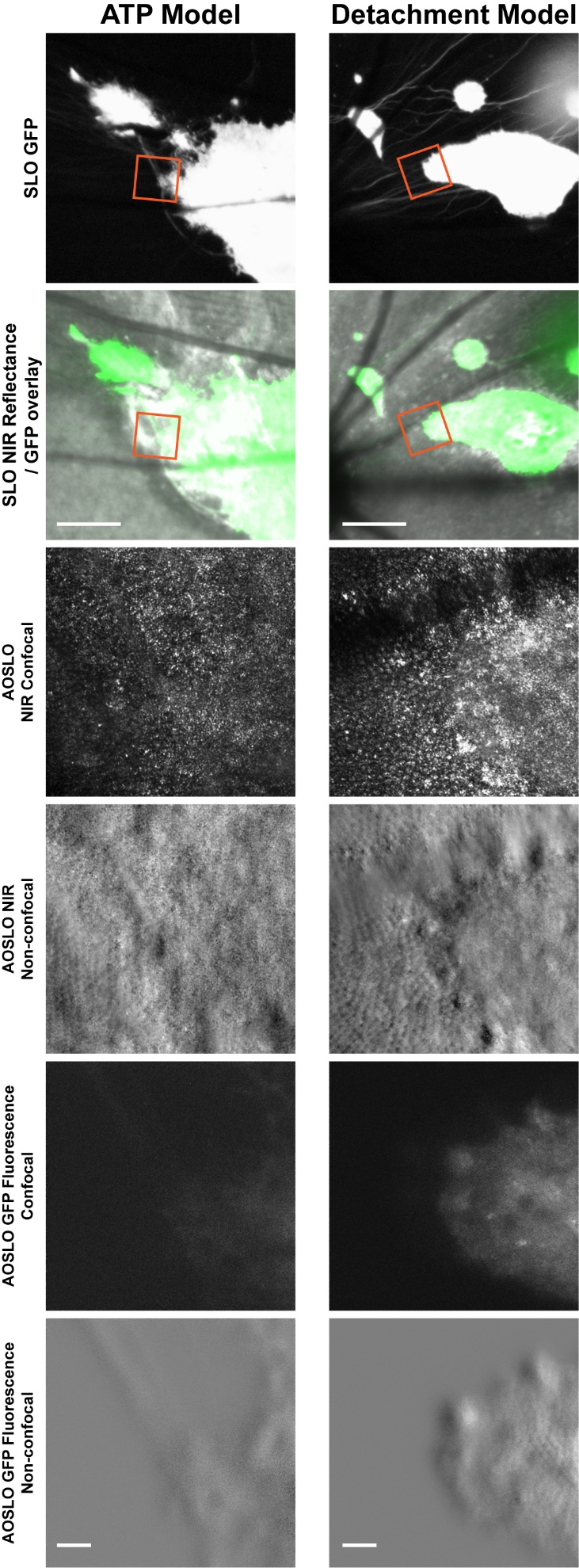

**Supplementary Figure S6. Fluorescence AOSLO imaging of transplanted tree shrew retina.** Fluorescence AOSLO was collected at the same location as NIR AOSLO in both confocal and non-confocal modality. NIR AOSLO captures the host cone photoreceptors, and fluorescence AOSLO captures the transplanted cone photoreceptors. Scale bars are 5 degrees (SLO images), 30μm (AOSLO images).
