## Supplementary material for "Transplantation of human iPSC-derived retinal cells in the cone-dominant Northern tree shrew": Figure S7

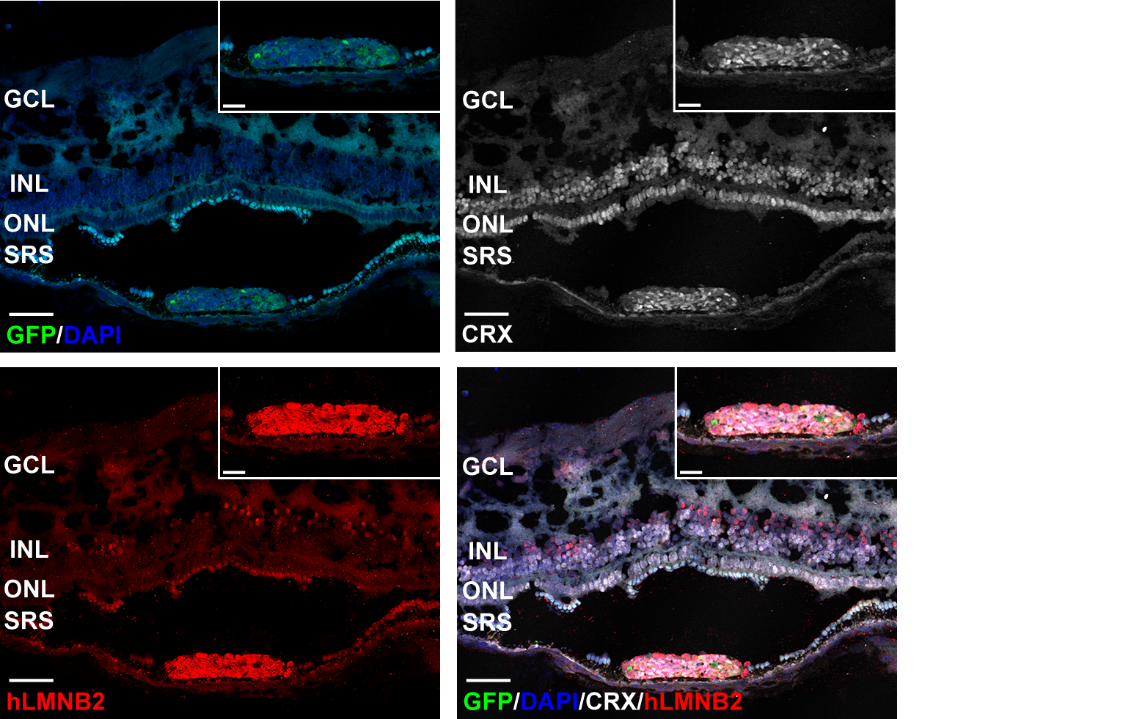

**Supplementary Figure S7. Histological analysis of transplanted ATP damaged tree shrew eyes.** Representative immunofluorescence images of the ATP damaged eye with transplanted graft showing the location of the GFP+ cells, human marker (hLMNB2) expression at the transplant location, photoreceptor cell (CRX) expression across the retina, merge of all markers. Scale bars are 50 μm (large image) and 20 μm (inset image). SRS, subretinal space; ONL, outer nuclear layer; INL, inner nuclear layer; GCL, ganglion cell layer.
