## Supplementary material for "Transplantation of human iPSC-derived retinal cells in the cone-dominant Northern tree shrew": Table S1

**Table S1**. Animal damage method and transplanted dates

| **Eye Number** | **Type of Damage** | **Animal ID_Eye** | **D.O.B.** | **Sex** | **Damage Notes (ATP Dose or Detachment Saline Volume)** | **Date of Damage Creation** | **Weight (Date of iPSC injection)** | **Date of iPSC Injection** |
| --- | --- | --- | --- | --- | --- | --- | --- | --- |
| 1 | Detachment | 7DE29F2_OS | 07/24/18 | F | 80 µL saline | 9/22/21 | 267g | 2/23/22 |
| 2 | Detachment | 7D642BC_OD | 05/18/18 | M | 100 µL saline | 12/15/21 | 193g |  |
| 3 | Detachment | 7D642BC_OS |  |  | 100 µL saline | 12/15/21 |  |  |
| 4 | Detachment | 7A4E459_OD | 02/25/21 | M | 30 µL saline | 7/26/22 | 285g | 9/8/22 |
| 5 | None | 7DE2337_OD | 08/04/18 | F | N/A | N/A | 185g |  |
| 6 | None | 7DE2337_OS |  |  | N/A | N/A |  |  |
| 7 | None | 7A4BB96_OD | 01/27/21 | M | N/A | N/A | 184g | 11/2/22 |
| 8 | None | 7A4BB96_OS |  |  | N/A | N/A |  |  |
| 9 | None | 7A4C8DA_OD | 10/10/20 | F | N/A | N/A | 171g |  |
| 10 | Detachment | 7A4C8DA_OS |  |  | 30 µL saline | 7/26/22 |  |  |
| 11 | None | 7A4D956_OD | 10/10/20 | F | N/A | N/A | 150g | 1/10/23 |
| 12 | None | 7A4D956_OS |  |  | N/A | N/A |  |  |
| 13 | Detachment | 7DE21D2_OD | 08/08/18 | F | 30 µL saline | 12/8/22 | 122g |  |
| 14 | Detachment | 7DE21D2_OS |  |  | 30 µL saline | 12/8/22 |  |  |
| 15 | Detachment | 80F0D53_OS | 09/26/21 | M | 30 µL saline | 7/31/23 | 208g | 8/16/23 |
| 16 | ATP | 80F0D53_OD |  |  | 0.01 M ATP | 6/1/23 |  |  |
| 17 | Detachment | 7E6CA55_OS | 04/09/22 | M | 25-30 µL saline | 7/31/23 | 156g |  |
| 18 | ATP | 7E6CA55_OD |  |  | (1) 0.0075 M ATP  (2) 0.0085 M ATP | 6/12/23 |  |  |
| 19 | Detachment | 80F6566_OS | 03/03/22 | M | 30 µL saline | 7/31/23 | 202g | 8/22/23 |
| 20 | ATP | 80F6566_OD |  |  | (1) 0.0075 M ATP (2) 0.0085 M ATP | 6/12/23 6/26/23 |  |  |
| 21 | Detachment | 7E9FAAD_OS | 02/25/22 | M | 25-30 µL saline | 7/31/23 | 138g |  |
| 22 | ATP | 7E9FAAD_OD |  |  | 0.0085 M ATP | 7/21/23 |  |  |
