## Supplementary material for "Transplantation of human iPSC-derived retinal cells in the cone-dominant Northern tree shrew": Table S2

**Table S2.** Cell colocalization quantification (**related to Figure S3**)

| **Retinal layer counted** | **ONL/SRS** | **INL/GCL** | |
| --- | --- | --- | --- |
| **Marker** | **OTX2/HuNu** | **HuCD/HuNu** | **Brn3a/HuNu** |
| **Undamaged-dataset 1** | 49.7% | No HuNu+ cell | No HuNu+ cell |
| **Undamaged-dataset 2** | 54.1% | 85% | 46.9% |
| **Detachment** | 60.5% | 100% | 33.3% |
