## Supplementary material for "Transplantation of human iPSC-derived retinal cells in the cone-dominant Northern tree shrew": Table S3

**Table S3**. Primary Antibody List **(related to Figures 4, 5)**.

| **Protein Name** | **Host** | **Dilution** | **Source** | **Cat. No.** |
| --- | --- | --- | --- | --- |
| Brn-3A | Mouse | 1:100 | Santa Cruz Biotechnology, Inc. | sc-8429 |
| GFP | Rabbit | 1:500 | GeneTex | GTX113617 |
| HuC/D | Mouse | 1:100 | Life Technologies (Invitrogen) | A-21271 |
| HuNu | Mouse | 1:500 | Millipore Sigma | MAB1281 |
| Lamin B2 | Mouse | 1:500 | GeneTex | GTX628803 |
| Otx2 | Goat | 1:400 | R&D Systems | BAF1979 |
| Recoverin | Rabbit | 1:1000 | EMD Millipore | AB5585 |

**Abbreviations:** Brn-3A, brain-specific homeobox/POU domain protein 3A; GFP, green fluorescent protein; HuC/HuD, Hu-antigen D; HuNu, human nuclei; Otx2, Orthodenticle homolog 2.
