## Supplementary material for "Transplantation of human iPSC-derived retinal cells in the cone-dominant Northern tree shrew": Table S4

**Table S4**. Secondary Antibody List **(related to Figures 4, 5)**.

| **Species** | **Target** | **Conjugate** | **Dilution** | **Source** | **Cat. No.** |
| --- | --- | --- | --- | --- | --- |
| Donkey | Anti-mouse | Alexa Fluor 488 | 1:250 | Invitrogen | A-21202 |
| Donkey | Anti-mouse | Alexa Fluor 555 | 1:250 | Invitrogen | A-31570 |
| Donkey | Anti-mouse | Alexa Fluor 647 | 1:250 | Invitrogen | A-31571 |
| Donkey | Anti-rabbit | Alexa Fluor 488 | 1:250 | Invitrogen | A-21206 |
| Donkey | Anti-rabbit | Alexa Fluor 555 | 1:250 | Invitrogen | A-31572 |
| Donkey | Anti-rabbit | Alexa Fluor 647 | 1:250 | Invitrogen | A-31573 |
| Donkey | Anti-goat | Alexa Fluor 488 | 1:250 | Invitrogen | A-11055 |
| Donkey | Anti-goat | Alexa Fluor 555 | 1:250 | Invitrogen | A-21432 |
| Donkey | Anti-goat | Alexa Fluor 647 | 1:250 | Invitrogen | A-21447 |
| Goat | Anti-mouse | Alexa Fluor™ 555 | 1:250 | Invitrogen | A-21127 |
| Goat | Anti-mouse | Alexa Fluor™ 647 | 1:250 | Invitrogen | A-21242 |
